# Molecular evolution of protein sequences and codon usage in monkeypox viruses

**DOI:** 10.1101/2022.12.23.521708

**Authors:** Ke-jia Shan, Changcheng Wu, Xiaolu Tang, Roujian Lu, Wenjie Tan, Jian Lu

## Abstract

The monkeypox virus (mpox virus, MPXV) epidemic in 2022 poses a significant public health risk. However, the evolutionary principles of MPXV are largely unknown. Here, we examined the evolutionary patterns of protein sequences and codon usage in MPXV. We first showed the signal of positive selection in *OPG027* specifically in Clade I. We then discovered accelerated protein sequence evolution over time in 2022 outbreak-causing variants. We also found strong epistasis between amino acid substitutions located in different genes. Codon adaptation index (CAI) analysis revealed that MPXV tended to use more unpreferred codons than human genes and that the CAI decreased with time and diverged between clades, with Clade I > IIa and IIb-A > IIb-B. Although the decrease in fatality rate in the three groups matched the CAI pattern, it is unclear whether this is a coincidence or if the deoptimization of codon usage in MPXV induced a decrease in fatality. This study sheds new light on the processes influencing the evolution of MPXV in human populations.

## Introduction

The monkeypox virus (mpox virus, MPXV) epidemic in 2022 has caused substantial public health risks. MPXV is a linear double-stranded DNA virus that belongs to the *Poxviridae* family, *Chordopoxvirinae* subfamily, and *Orthopoxvirus* genus^1–3^. The genome of MPXV is approximately 197 kb in length and encodes approximately two hundred genes^4^. MPXV can infect multiple animal species, including humans, nonhuman primates, and rodents^5–8^. MPXV, like variola virus (VARV) and vaccinia virus (VACV) in the *Orthopoxvirus* genus, can cause disease and death in humans.

In 1958, MPXV was discovered in a Danish animal facility^8^; in 1970, it was isolated from the first human case in the Democratic Republic of the Congo^9^. Prior to 2022, MPXV was predominantly endemic in the countries of Central and Western Africa, with spontaneously reported instances in other regions resulting from importations^10–15^. On May 7, 2022, the first human case of MPXV in the outbreak of 2022 was reported in the United Kingdom^16,17^. The global outbreak of MPXV was confirmed as an international public health emergency on July 23, 2022^18^. As of November 13, 2022, 79,411 confirmed cases had been recorded from 109 countries and regions in the 2022 outbreak^19^.

MPXV was classified into two clades based on phylogenetic analysis^20^: Clade I (also known as the ‘Central African’ clade) and Clade II (also known as the ‘West African’ clade). Clade II was further subdivided into the IIa and IIb subclades. The MPXV variants collected in the 2017-2018 outbreak corresponded to subclade IIa or lineage A in IIb (IIb-A)^21,22^. The majority of the MPXV variants that triggered the 2022 outbreak belong to IIb-B, and they are phylogenetically more closely related to the variants transported from Nigeria to the United Kingdom, Israel, and Singapore in 2018-2019 than to the variants collected during the 2017-2018 Nigeria outbreak^17,20^. In the 2022 outbreak, however, sporadic cases of IIb-A.2 sublineages have also been reported^17,20^.

The mutation rate of orthopoxviruses is 1-2 substitutions per genome per year^23^. Nevertheless, the evolutionary analysis revealed approximately 50 nucleotide differences in the 2022 variants compared to the 2018-2019 variants, which was 6-12 times greater than expected based on the usual orthopoxvirus mutation rate^17^. Moreover, excessive TC>TT and GA>AA mutations have been identified between the two clades of variants. It was hypothesized that the increased substitution rate in the 2022 MPXV genomes is the result of genome editing by apolipoprotein B mRNA-editing catalytic polypeptide-like 3 (APOBEC3) enzymes, which cause C>T mutations if editing occurs in the sense strand and G>A mutations if editing occurs in the antisense strand^17,20^. Since APOBEC3 enzymes are primate-specific^24^, it is plausible that the recent circulation of MPXV in humans has resulted in a higher mutation rate due to APOBEC3 editing than in its natural animal hosts, which include numerous non-primate species^20^.

Despite these recent discoveries, the evolutionary principles of MPXV remain poorly understood. Here, we first identified signatures of positive selection on MPXV genes. Then, we investigated epistasis in 2022 outbreak-causing MPXV variants. Finally, we examined MPXV codon usage patterns and their potential impact on fatality rates. This study revealed the accelerated evolution of the 2022 outbreak-causing MPXV variants and cautions the possible relationship between deoptimization codon usage and the decreased fatality rate in the MPXV evolutionary process.

## Results

### Positive selection on *OPG027* in Clade I of MPXV

To detect signatures of positive selection, we downloaded 1,953 high-quality MPXV genome sequences from the National Center for Biotechnology Information (NCBI)^25^ and GISAID^26^ (https://www.gisaid.org, as of November 13, 2022) and divided these sequences into four clades (I, IIa, IIb-A, and IIb-B) and calculated the dN (nonsynonymous substitutions per nonsynonymous site), dS (synonymous substitutions per synonymous site), and dN/dS (ω) values for each gene between any two genomes from different clades. The majority of the comparisons yielded ω < 1, showing that purifying selection is the dominant force driving MPXV gene evolution (**Fig. 1a**). Interestingly, we found ω >1 for *OPG027* in pairwise comparisons between clades I and II, suggesting that this gene was subjected to positive selection during the divergence of these two clades. However, *OPG027* showed strong signatures of purifying selection (ω < 1) in the pairwise comparisons between Clade II genomes. To better understand the discrepancies between these findings, we analyzed the SNPs in *OPG027* across the 1,953 MPXV genomes, using VACV and VARV as outgroups. In the branch from the most recent common ancestor of Clades I and II to Clade I, we found three nonsynonymous changes (I47V, F79L, and V126I) and one synonymous nucleotide substitution (C276T). In contrast, no substitutions were found on the Clade II branch (**Fig. 1b** and **Table S1**). These data imply that *OPG027* has experienced accelerated protein sequence evolution specifically in Clade I. OPG027 is crucial in determining host ranges and inhibiting host antiviral activities^25,27,28^. The accelerated evolution of *OPG027* in Clade I could have been triggered by evolutionary arms races related to viral replication or immune evasion.

**Figure 1.**
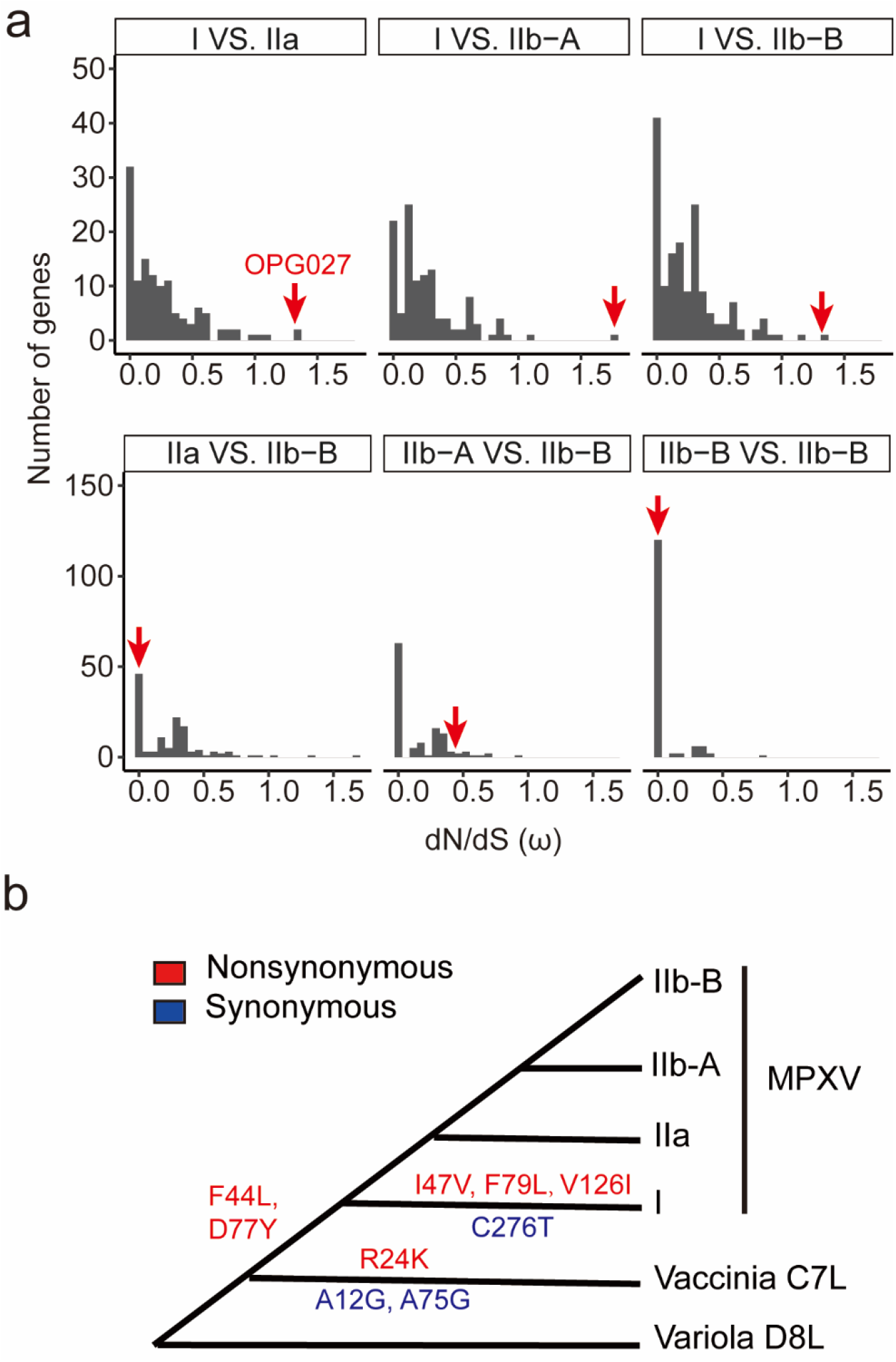
Positive selection on *OPG027*. (**a**) The distribution of the median dN/dS ratio (ω) between two sequences of each MPXV gene from different clades. The ω values of *OPG027* are indicated by the red arrows. (**b**) The amino acid substitutions of OPG027 in the evolutionary process of MPXV.

### Accelerated protein evolution in 2022 outbreak-causing MPXV variants

To decipher the evolutionary trends of currently circulating MPXV variants, we analyzed 756 IIb-B genomes with exact collection dates throughout the 2022 outbreak. In one MPXV genome, the median number of substitutions relative to the reference genome (NC_063383, collected in August 2018 in Rivers State, Nigeria) was 69 (2.5th and 97.5th percentiles of 67 and 81, respectively), while the median numbers of synonymous and nonsynonymous SNPs were 27 (2.5th and 97.5th percentiles of 27 and 30, respectively) and 33 (the 2.5th and 97.5th percentiles of 32 and 41, respectively).

TreeTime^29^ analysis of these 756 MPXV genomes yielded a substitution rate of 6.07 ± 0.86 × 10^-5^ substitutions/site/year at the genomic scale and lower substitution rates in the coding regions (the substitution rates in the first, second, and third positions of the codons were 5.70 ± 2.70 × 10^-5^, 3.17 ± 2.52 × 10^-5^ and 4.66 ± 2.56 × 10^-5^ substitutions/site/year, respectively). Because nucleotide changes in the second position of a codon are nonsynonymous, while changes in the third position of a codon are mostly synonymous, these findings support the notion that purifying selection is the dominant evolutionary force affecting MPXV protein sequences.

The number of substitutions in an MPXV genome increased with the time between the sample’s collection date and the reference genome availability date (which was arbitrarily set on August 15, 2018) **(Fig. 2a-c)**. Notably, the number of nonsynonymous substitutions showed a significantly higher correlation coefficient with time than the number of synonymous substitutions (*Rho* was 0.33 versus 0.16 for the former versus the latter, respectively; *P* = 0.0008, Fisher’s method). Remarkably, the slope of the linear regression between nonsynonymous substitutions and time was significantly steeper than that of the linear regression between synonymous substitutions and time (the slopes were 0.029 versus 0.0046, respectively; *P* = 0.001). These findings support the notion that the 2022 outbreak-causing MPXV variants have experienced accelerated protein sequence evolution.

**Figure 2.**
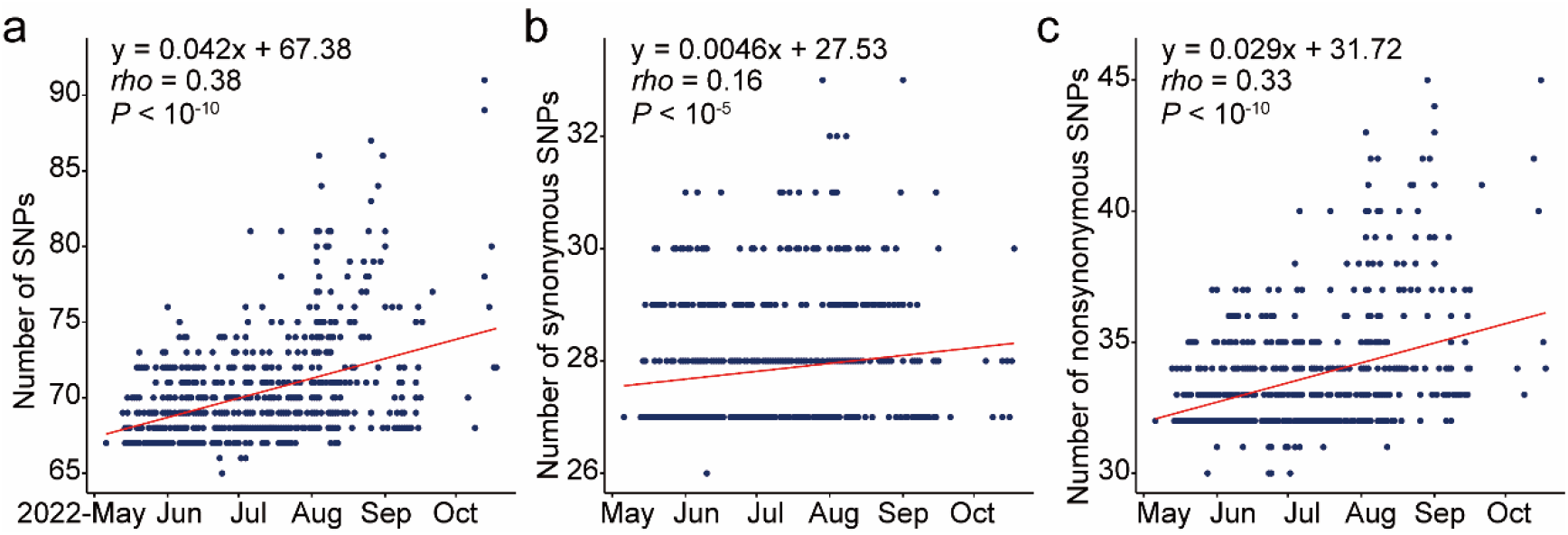
SNPs of lineage B during the 2022 monkeypox outbreak. The number of SNPs in the whole genome (**a**), at synonymous sites (**b**), and at nonsynonymous sites (**c**). The *y*-axis indicates the number of SNPs in each genome sequence relative to the reference genome (NC_063383). The *x*-axis indicates the collection date.

### Epistasis in 2022 outbreak-causing MPXV variants

We analyzed the patterns of linkage disequilibrium (LD) between MPXV SNPs (**Fig. S1**). We focused on SNPs with frequencies ranging from 0.005 to 0.8 in the 1,873 MPXV genomes of Clade IIb-B, and only SNP pairs with *r^2^* ≥ 0.8 were considered in the LD analysis. Under these criteria, we discovered 41 substitutions (16 synonymous, 21 nonsynonymous, and 4 intergenic ones) forming 15 linkage groups that were distributed in 12 sublineages of Clade IIb-B (**Fig. 3** and **Table S2**). Two linkage groups (Groups 6 and 8) had just synonymous or noncoding alterations, and six groups (Groups 1, 2, 3, 4, 12, and 15) had one nonsynonymous substitution plus at least one synonymous (or noncoding) substitution (**Table S2**). Notably, six linkage groups (Groups 5, 7, 9, 10, 11, and 13) were composed of at least two tightly linked nonsynonymous substitutions located in different genes, such as H173Y in *OPG038* (NF-κB inhibitor) and D124N in *OPG099* (Membrane protein CL5) in the case of B.1.8, D162N in *OPG040* (Serpin) and R88K in *OPG107* (Entry-fusion complex essential component) in the case of B.1.4, and S288L in *OPG185* (Hemagglutinin) and S156L in *OPG055* (Protein F11) in the case of B.1.14. One linkage group, in particular, consisted of one synonymous substitution (V273V in *OPG130*) and three nonsynonymous substitutions: S532L in *OPG210* (B22R family protein), D729N in *OPG117* (NTPase), and G4R in *OPG118* (Early transcription factor 70 kDa subunit) (**Table S2**). We speculated that these epistatic interactions would be associated with the fitness of an MPXV lineage because many genes with these tightly linked amino acid alterations are involved in viral infection or anti-host immunity. More functional research is needed to determine the biological functions of these changes as well as their epistatic impacts.

**Figure 3.**
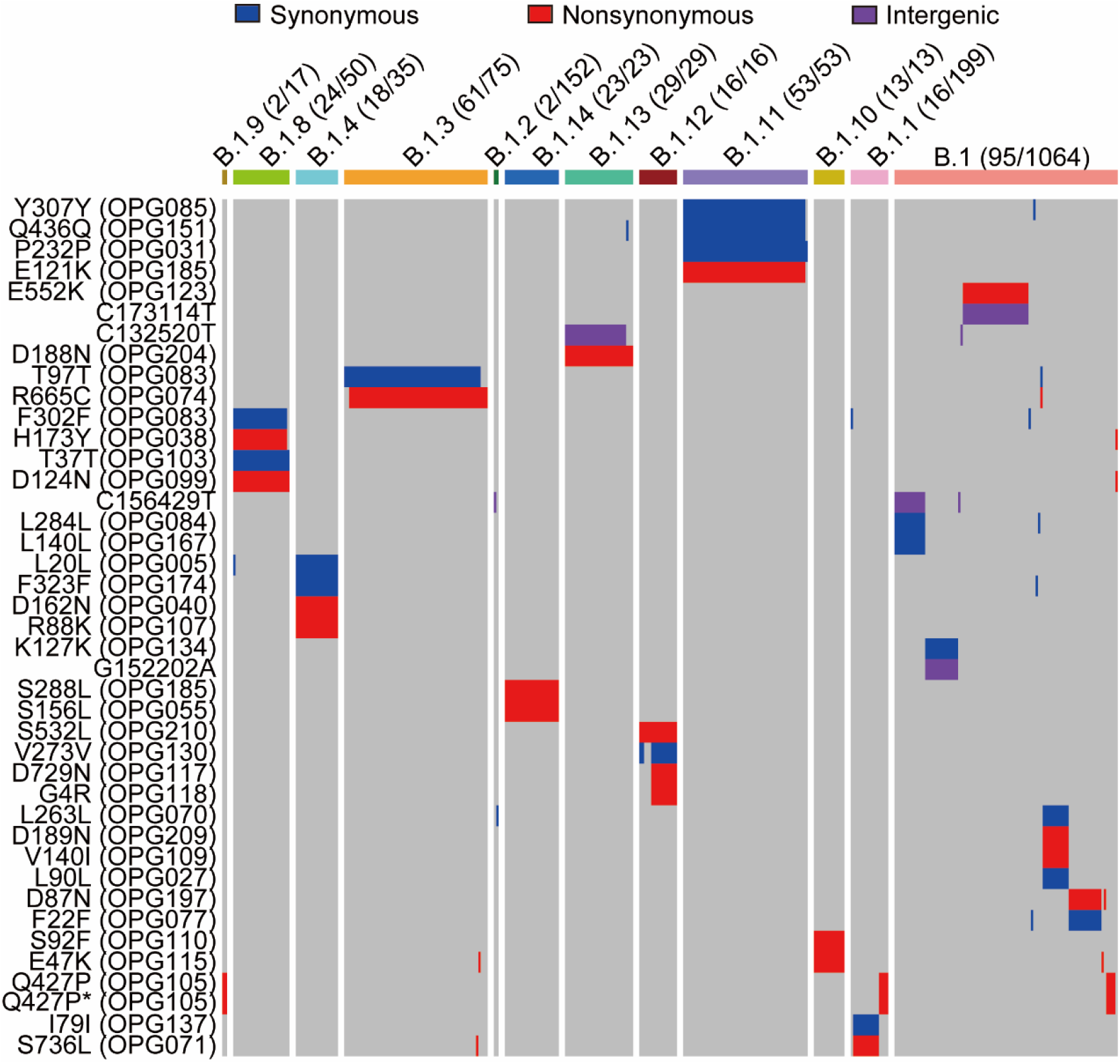
Linkage disequilibrium between SNPs in Clade IIb-B of MPXV. The fifteen linked SNP groups (*r^2^* ≥ 0.8) in Clade IIb-B of MPXV. The *y*-axis indicates the amino acid substitutions in the proteins caused by SNPs or SNPs in the intergenic region. The A82405C/A82406T causing CAA (Q) > CCT (P) in *OPG105* is labeled Q427P/Q427P*. The number of sequences containing at least one of these 41 substitutions and the number of sequences used to detected LD are shown as the numerator and denominator, respectively.

### Deoptimization of codon usage in MPXV over time

Viruses often rely on the host organism’s cellular machinery for biological functions such as translation. They also often exhibit a low level of codon use bias, owing to mutational pressure or natural selection^30,31^. Viruses with poor codon usage are proposed to be more adaptable to different host species^31–33^.

To examine the codon usage bias in MPXV variants, we calculated the codon adaptation index (CAI) of the concatenated coding sequences (CDSs) in each MPXV genome as previously described^34^. The MPXV CAI values varied from 0.6093 to 0.6104, with 0.6098 as the median and 0.6098 and 0.6100 as the 2.5th and 97.5th percentiles, respectively. Overall, the CAI value of MPXV was substantially lower than those of human genes (**Fig. 4a**), suggesting that MPXV is more likely to use unpreferred codons than human genes. This observation is in line with the notion that MPXV genomes are A/T rich because codons rich in A/T nucleotides are generally unpreferred in humans.

**Figure 4.**
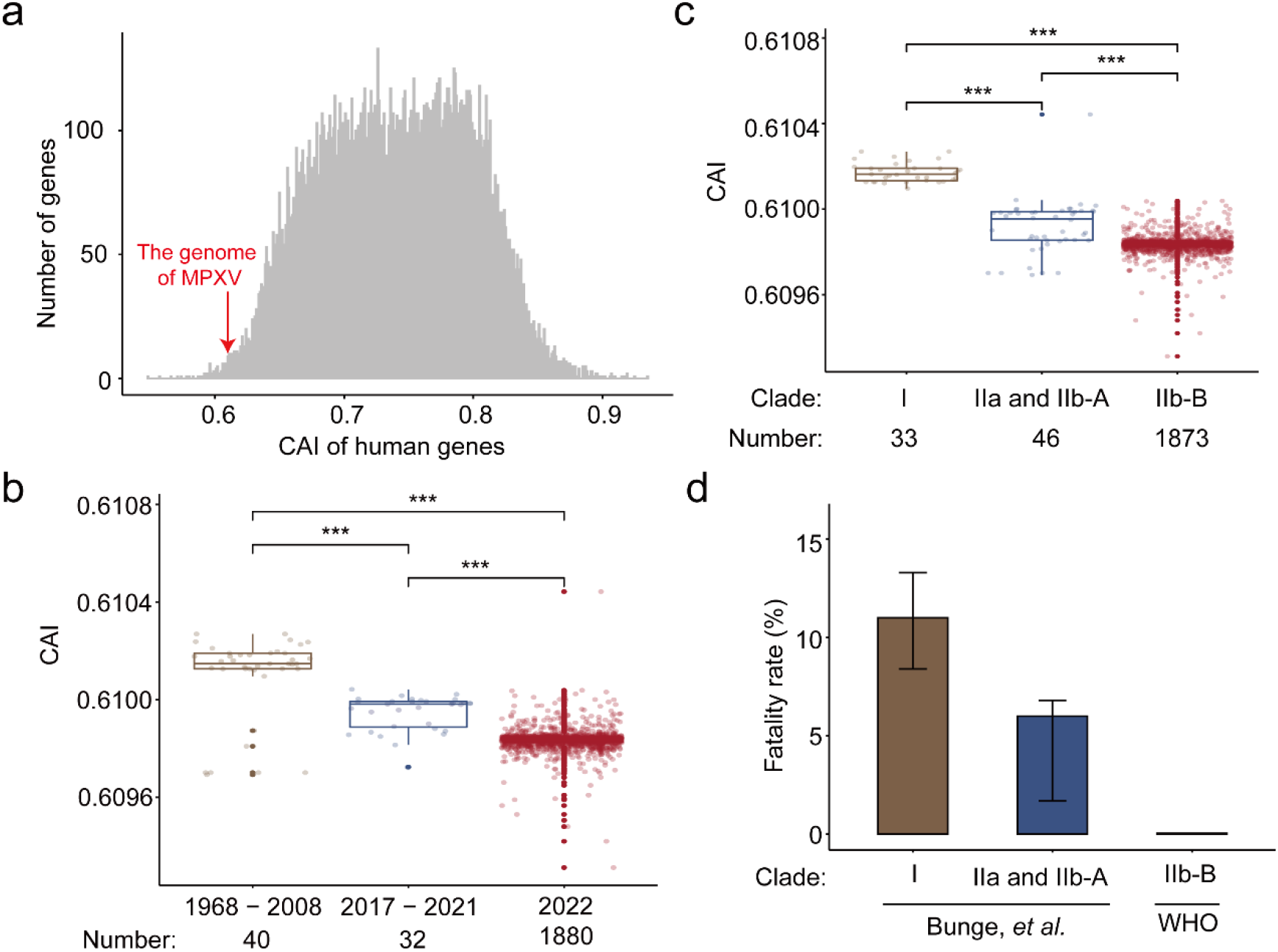
The CAI and fatality rate of MPXV. (**a**) The distribution of CAI values of human genes and the concatenated coding sequences of MPXV. (**b**) The CAI values decreased over time. (**c**) The CAI of different clades and lineages. (**d**) The fatality rate of different clades and lineages. The error bars indicate the 95% confidence intervals.

When we grouped the MPXV variants into three categories based on the collection dates in 1968-2008, 2017-2021, and 2022, we found that the CAI of MPXV decreased over time, and this trend was statistically significant (**Fig. 4b**). Because Clade I is the oldest and Clade IIb-B is the most recent, it is not surprising that a significant difference in the CAI was found when we separated the MPXV genomes into three groups based on lineage, in the order of Clade I > IIa and IIb-A > IIb-B (**Fig. 4c**). The continued deoptimization of codons in MPXV genomes was most likely caused by an overabundance of C>T or G>A mutations driven by APOBEC3-mediated viral editing **(Table S3)**.

MPXV Clade I had a fatality rate of 10.6% (95% CI: 8.4–13.3%), while clades IIa and IIb-A had a fatality rate of 3.6% (95% CI: 1.7%–6.8%)^10^. Clade IIb-B, which primarily caused the 2022 MPXV outbreak, had a fatality rate of 0.063% (50 deaths out of 79,411 confirmed cases, according to the WHO as of November 13, 2022)^19^. Notably, there was a significant difference in fatality rate among the MPXV clades, with Clade I > IIa and IIb-A > IIb-B (**Fig. 4d**). Although the decline in fatality rate followed a pattern similar to that of CAI in the three groups, it is unclear if this is simply a coincidence or the consequence of a causal relationship in which the deoptimization of codon usage caused a decrease in fatality across the three groups.

## Discussion

The pathogenicity and drug resistance of viruses could be significantly boosted by only a few amino acid changes. For instance, an amino acid substitution (N752D) in the DNA polymerase of equid herpesvirus type 1 (EHV-1) could significantly alter its neuropathogenicity^35^, and one amino acid change (T831I) or two linked changes (A314V and A684V) in the DNA polymerase (E9L) of VACV significantly increased levels of drug resistance^36^. In this study, we detected signals of positive selection in *OPG027* specifically in Clade I of MPXV. Because OPG027 is important in determining host range and inhibiting type-1 interferon^27,28^, the amino acid changes (I47V, F79L, and V126I) in *OPG027* may serve as candidates for future functional studies to investigate the biological difference between Clade I and II variants.

Similar to findings for SARS-CoV-2^37–39^, we detected many tightly linked amino acid changes in the 2022 outbreak-causing MPXV variants. These changes tend to be located in different genes, most of which are associated with viral entry or immune evasion. It is plausible that compensatory advantageous mutations occurred during the 2022 outbreak, which could explain the accelerated protein sequence evolution in these MPXV variants. However, we cannot rule out the possibility that sampling bias or founder effects influenced the observed trends. Future research should examine the evolutionary driving mechanisms and biological significance of these epistatic interactions.

Codon usage bias can affect protein expression and function by changing translation efficiency^40^, mRNA stability^41^, and peptide conformation^42,43^. MPXV, similar to SARS-CoV-2 and many other viruses^34^, tends to use more unpreferred codons than human genes. The CAI of MPXV also declined with time and differed between clades, with Clade I > IIa and IIb-A > IIb-B. Notably, the fatality rate also differed significantly among the MPXV clades, in the order of Clade I > IIa and IIb-A > IIb-B. Since viruses with significant translational activities may impose a translational burden on the host or cause severe clinical symptoms^44^, the deoptimization of codon usage in MPXV might cause the virus to replicate slowly and thus reduce the fatality rate during evolution. However, we cannot rule out the possibility that the similar trends in the CAI and fatality rates are simply coincidences caused by sampling bias or other confounding factors. Further studies are needed to decipher the observed relationship. There is a need for vaccine development to combat MPXV^45^, and our findings show the importance of optimizing codon usage in mRNA and DNA vaccine design because codon usage impacts the efficiency of antigen expression.

## Materials and methods

### Evolutionary analysis

A total of 2,789 MPXV genome sequences were retrieved from the National Center for Biotechnology Information (NCBI)^25^ and GISAID^26^ (https://www.gisaid.org, as of November 13, 2022). Only 1,953 complete and unique sequences were used for downstream analysis. Multiple sequence alignment and mutation identification were performed by Nextclade v2.4.0^46^. The mutations were annotated by SnpEff v5.0e^47^ based on the reference genome (NCBI: NC_063383). Only 756 IIb-B genomes with exact collection dates were used to estimate the mutation rates based on the phylogenetic relationships by TreeTime v0.9.4 (--reroot oldest --covariation) ^29^.

### The detection of positive selection

For each gene, we kept only one sequence of pairs of identical sequences and discarded sequences containing more than 15 ambiguous nucleotides or gaps. Then, we calculated N (the number of nonsynonymous sites), S (the number of nonsynonymous sites), dN (nonsynonymous mutations per nonsynonymous site), dS (synonymous mutations per synonymous site) and the dN/dS (ω) ratio of every sequence pair by implementing the yn00 program in PAML v4^48^.

### The linkage disequilibrium of Clade IIb-B

We calculated the *r^2^* (square of the correlation coefficient) of each SNP pair outside the inverted terminal repeat regions of Clade IIb-B using an in-house script. Each SNP was supported by at least five genome sequences and had a frequency of less than 0.8 but more than 0.005 in Clade IIb-B. Only the SNP pairs with *r^2^* ≥ 0.8 were selected as linked SNPs.

### Calculation of the codon adaptation index (CAI) of MPXV

The CAI was calculated as previously described^34^. In brief, we weighted the frequencies of codons based on the median expression levels in 54 human tissues from the Genotype-Tissue Expression (GTEx) database Version 8 (https://www.gtexportal.org/). Then, the CAI was calculated according to the actual frequencies of codons in the transcriptomes.

We extracted the coding sequences of each MPXV sequence based on the multiple sequence alignment from Nextclade^46^ and the annotation of the reference genome (NCBI: NC_063383). We concatenated the coding sequences of MPXV to calculate the human-expression weighted CAI value.

## Acknowledgments

We thank the researchers who generated and shared the sequencing data in the NCBI and GISAID (https://www.gisaid.org/) databases, on which this research is based. This work was supported by the National Natural Science Foundation of China (82241080), the National Key Research and Development Projects of the Ministry of Science and Technology of China (2021YFC2301300, 2022YFC2304100, 2022YFC2303401), and the SLS-Qidong Innovation Fund.

## Supplementary Tables and Figures

**Table S1.** The number of sequences with mutations in *OPG027* among different clades of MPXV.

| SNP | Type<sup>a</sup> | AA change | Clade |  |  |  |
| --- | --- | --- | --- | --- | --- | --- |
|  |  |  | I | IIa | IIb-A | IIb-B |
| G16333A | syn | I15I | - | - | - | 1/1873 |
| C16327T | syn | L17L | - | - | - | 2/1873 |
| T16239C | nonsyn | I47V | 33/33 | - | - | - |
| C16227T | nonsyn | D51N | - | - | 1/31 | - |
| G16213A | syn | I55I | - | - | - | 1/1873 |
| C16209T | nonsyn | E57K | - | - | - | 1/1873 |
| G16147A | syn | Y77Y | 1/33 | - | - | - |
| G16141T | nonsyn | F79L | 32/33 | - | - | - |
| C16125T | nonsyn | E85K | - | - | - | 1/1873 |
| C16123A | nonsyn | E85D | 12/33 | - | - | - |
| G16110A | syn | L90L | - | - | - | 11/1873 |
| G16102A | syn | N92N | 33/33 | - | - | - |
| C16002T | nonsyn | V126I | 33/33 | - | - | - |
| C15947T | nonsyn | R144K | - | - | - | 1/1873 |

<sup>a</sup>syn: synonymous; nonsyn: nonsynonymous.

**Table S2.**
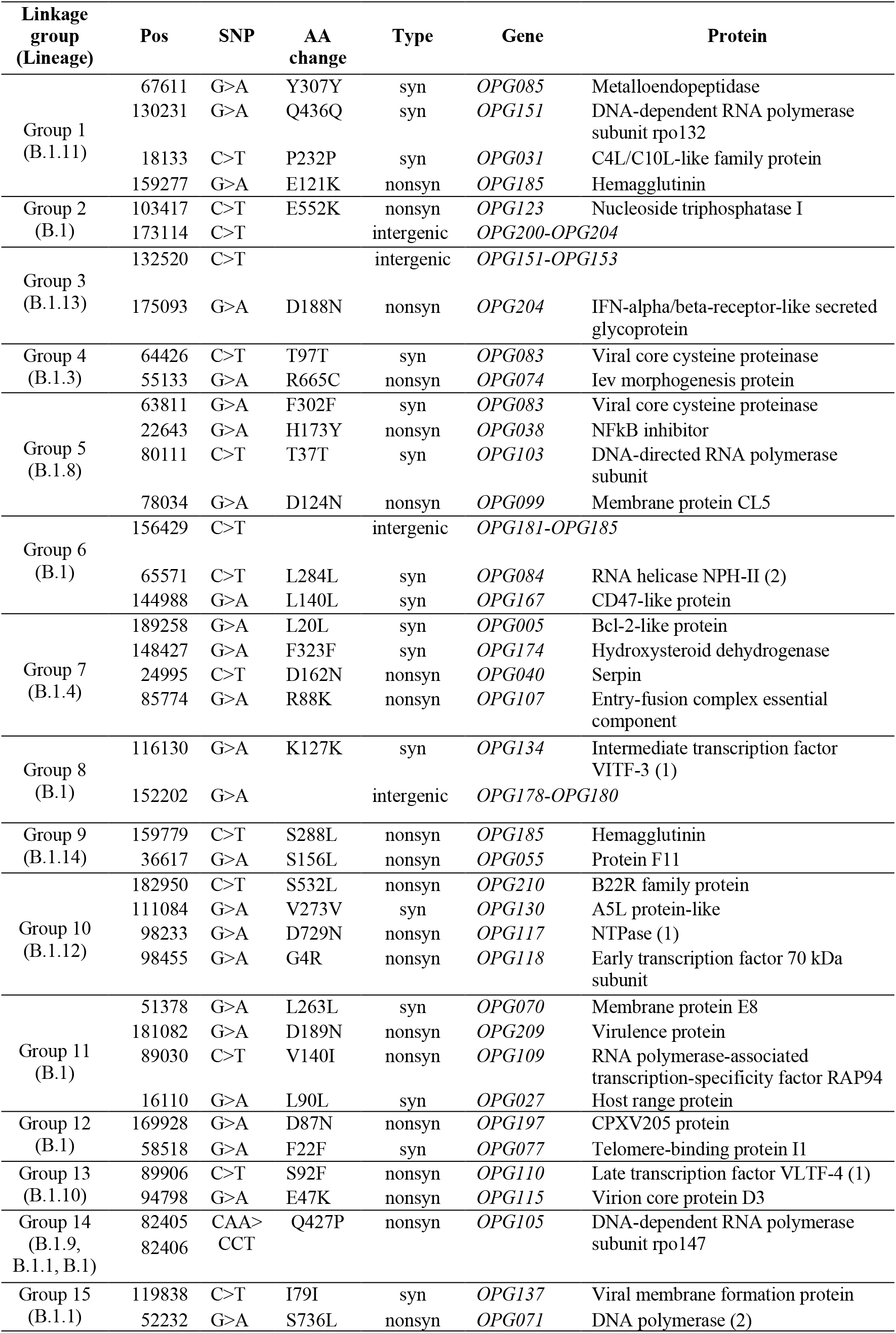
The linkage group of substitutions in Clade IIb-B of MPXV.

**Table S3.**
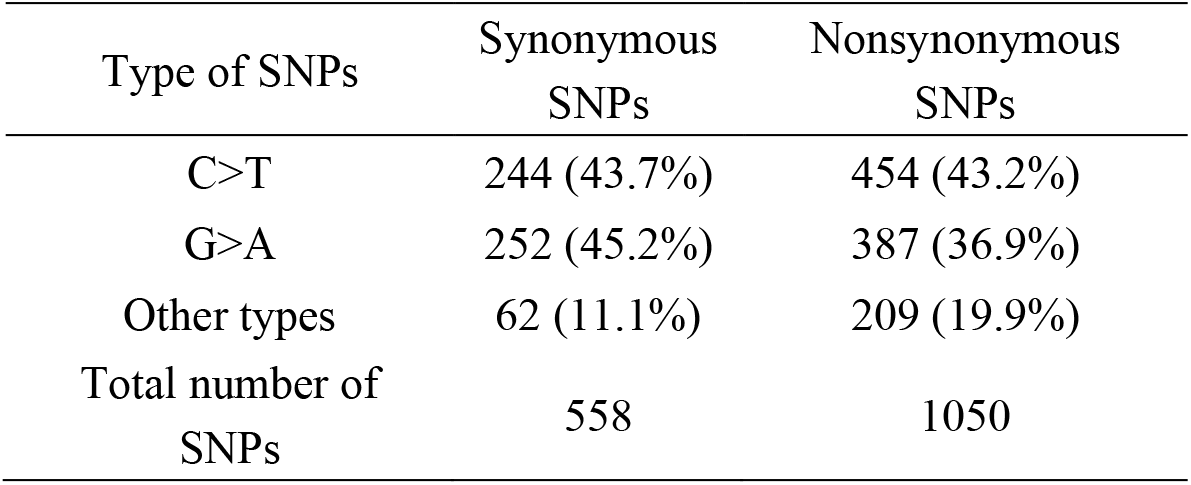
Summary of SNPs in Clade IIb-B of MPXV.

| Type of SNPs | Synonymous<br>SNPs | Nonsynonymous<br>SNPs |
| --- | --- | --- |
| C>T | 244 (43.7%) | 454 (43.2%) |
| G>A | 252 (45.2%) | 387 (36.9%) |
| Other types | 62 (11.1%) | 209 (19.9%) |
| Total number of<br>SNPs | 558 | 1050 |

**Figure S1.**
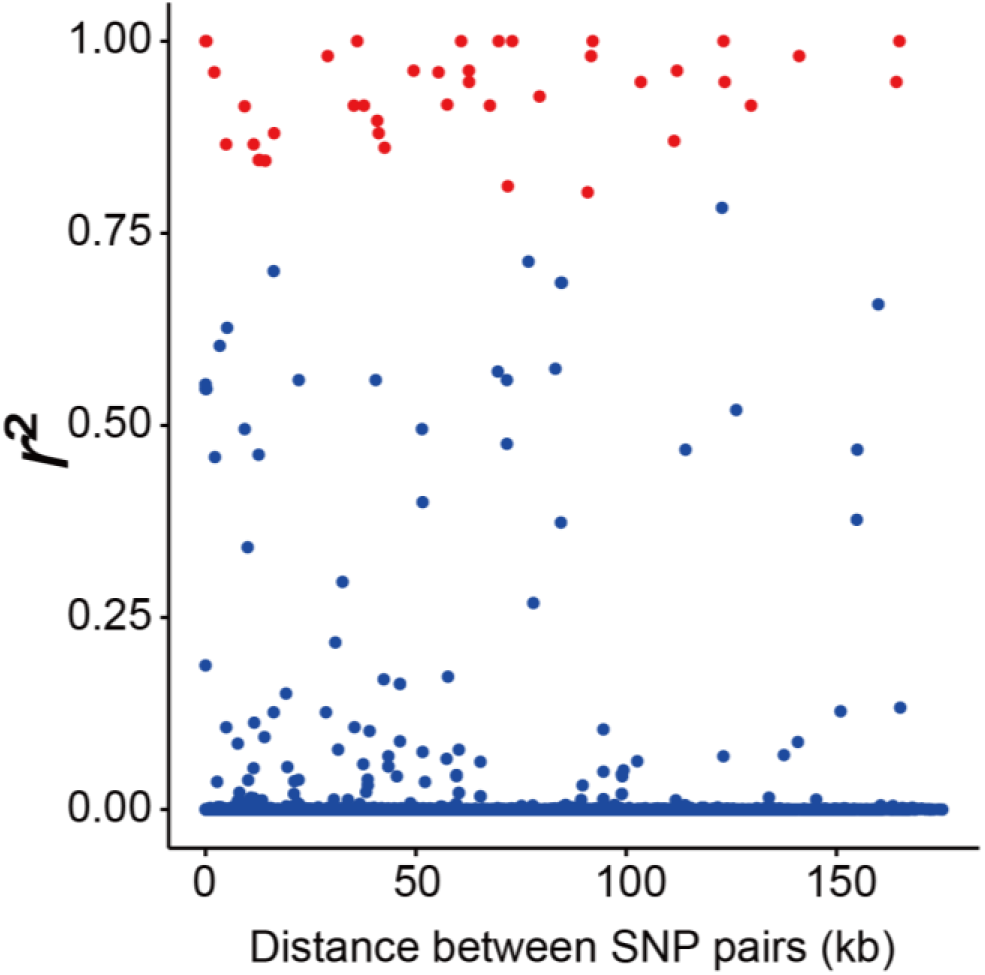
Linkage disequilibrium between SNPs in Clade IIb-B of MPXV. The *r^2^* (*y*-axis) against the distance between each SNP pair (*x*-axis). The SNP pairs in the fifteen linked SNP groups (*r^2^* ≥ 0.8) included in **Fig. 3** are shown as red points.

## References

1 Samaranayake, L. & Anil, S. The Monkeypox Outbreak and Implications for Dental Practice. Int Dent J 72, 589–596, doi:10.1016/j.identj.2022.07.006 (2022).

2 Petersen, E. et al. Human Monkeypox Epidemiologic and Clinical Characteristics, Diagnosis, and Prevention. Infect Dis Clin N Am 33, 1027–1043, doi:10.1016/j.idc.2019.03.001 (2019).

3 Seitz, R. et al. Orthopox Viruses: Infections in Humans. Transfus Med Hemoth 37, 351–364, doi:10.1159/000322101 (2010).

4 Shchelkunov, S. N. et al. Analysis of the monkeypox virus genome. Virology 297, 172–194, doi:10.1006/viro.2002.1446 (2002).

5 Doty, J. B. et al. Assessing Monkeypox Virus Prevalence in Small Mammals at the Human-Animal Interface in the Democratic Republic of the Congo. Viruses 9, 283, doi:10.3390/v9100283 (2017).

6 Radonic, A. et al. Fatal monkeypox in wild-living sooty mangabey, Cote d’Ivoire, 2012. Emerg Infect Dis 20, 1009–1011, doi:10.3201/eid2006.13-1329 (2014).

7 Haddad, N. The presumed receptivity and susceptibility to monkeypox of European animal species. Infect Dis Now 52, 294–298, doi:10.1016/j.idnow.2022.06.006 (2022).

8 Parker, S. & Buller, R. M. A review of experimental and natural infections of animals with monkeypox virus between 1958 and 2012. Future Virol 8, 129–157, doi:10.2217/fvl.12.130 (2013).

9 Ladnyj, I. D., Ziegler, P. & Kima, E. A human infection caused by monkeypox virus in Basankusu Territory, Democratic Republic of the Congo. Bull World Health Organ 46, 593–597 (1972).

10 Bunge, E. M. et al. The changing epidemiology of human monkeypox-A potential threat? A systematic review. PLoS Negl Trop Dis 16, e0010141, doi:10.1371/journal.pntd.0010141 (2022).

11 Centers for Disease, C. & Prevention. Update: multistate outbreak of monkeypox--Illinois, Indiana, Kansas, Missouri, Ohio, and Wisconsin, 2003. MMWR Morb Mortal Wkly Rep 52, 561–564 (2003).

12 Yinka-Ogunleye, A. et al. Outbreak of human monkeypox in Nigeria in 2017-18: a clinical and epidemiological report. Lancet Infect Dis 19, 872–879, doi:10.1016/S1473-3099(19)30294-4 (2019).

13 Mauldin, M. R. et al. Exportation of Monkeypox Virus From the African Continent. J Infect Dis 225, 1367–1376, doi:10.1093/infdis/jiaa559 (2022).

14 Costello, V. et al. Imported Monkeypox from International Traveler, Maryland, USA, 2021. Emerg Infect Dis 28, 1002–1005, doi:10.3201/eid2805.220292 (2022).

15 Rao, A. K. et al. Monkeypox in a Traveler Returning from Nigeria - Dallas, Texas, July 2021. MMWR Morb Mortal Wkly Rep 71, 509–516, doi:10.15585/mmwr.mm7114a1 (2022).

16 World Health Organization. Monkeypox - United Kingdom of Great Britain and Northern Ireland. https://www.who.int/emergencies/disease-outbreak-news/item/2022-DON2381 (2022).

17 Isidro, J. et al. Phylogenomic characterization and signs of microevolution in the 2022 multi-country outbreak of monkeypox virus. Nat Med 28, 1569–1572, doi:10.1038/s41591-022-01907-y (2022).

18 World Health Organization. WHO Director-General’s statement at the press conference following IHR Emergency Committee regarding the multi-country outbreak of monkeypox - 23 July 2022. https://www.who.int/director-general/speeches/detail/who-director-general-s-statement-on-the-press-conference-following-IHR-emergency-committee-regarding-the-multi--country-outbreak-of-monkeypox--23-july-2022 (2022).

19 World Health Organization. Multi-country outbreak of monkeypox. https://www.who.int/publications/m/item/multi-country-outbreak-of-monkeypox--external-situation-report--10---16-november-2022 (2022).

20 Gigante, C. M. et al. Multiple lineages of monkeypox virus detected in the United States, 2021–2022. Science 378, 560–565, doi:doi:10.1126/science.add4153 (2022).

21 Forni, D., Cagliani, R., Molteni, C., Clerici, M. & Sironi, M. Monkeypox virus: The changing facets of a zoonotic pathogen. Infect Genet Evol 105, 105372, doi:10.1016/j.meegid.2022.105372 (2022).

22 Luna, N. et al. Phylogenomic analysis of the monkeypox virus (MPXV) 2022 outbreak: Emergence of a novel viral lineage? Travel Med Infect Di 49, 102402, doi:10.1016/j.tmaid.2022.102402 (2022).

23 Firth, C. et al. Using time-structured data to estimate evolutionary rates of double-stranded DNA viruses. Mol Biol Evol 27, 2038–2051, doi:10.1093/molbev/msq088 (2010).

24 Jarmuz, A. et al. An anthropoid-specific locus of orphan C to U RNA-editing enzymes on chromosome 22. Genomics 79, 285–296, doi:10.1006/geno.2002.6718 (2002).

25 Sayers, E. W. et al. Database resources of the national center for biotechnology information. Nucleic Acids Res., D10–D17, doi:10.1093/nar/gkab1112 (2021).

26 Shu, Y. L. & McCauley, J. GISAID: Global initiative on sharing all influenza data - from vision to reality. Eurosurveillance 22, 2–4, doi:10.2807/1560-7917.Es.2017.22.13.30494 (2017).

27 Meng, X. Z. et al. C7L Family of Poxvirus Host Range Genes Inhibits Antiviral Activities Induced by Type I Interferons and Interferon Regulatory Factor 1. J Virol 86, 4538–4547, doi:10.1128/Jvi.06140-11 (2012).

28 Meng, X. Z. et al. Vaccinia Virus K1L and C7L Inhibit Antiviral Activities Induced by Type I Interferons. J Virol 83, 10627–10636, doi:10.1128/Jvi.01260-09 (2009).

29 Sagulenko, P., Puller, V. & Neher, R. A. TreeTime: Maximum-likelihood phylodynamic analysis. Virus Evol 4, vex042, doi:10.1093/ve/vex042 (2018).

30 Shackelton, L. A., Parrish, C. R. & Holmes, E. C. Evolutionary Basis of Codon Usage and Nucleotide Composition Bias in Vertebrate DNA Viruses. Journal of Molecular Evolution 62, 551–563, doi:10.1007/s00239-005-0221-1 (2006).

31 Jenkins, G. M. & Holmes, E. C. The extent of codon usage bias in human RNA viruses and its evolutionary origin. Virus Res 92, 1–7, doi:10.1016/s0168-1702(02)00309-x (2003).

32 Carmi, G., Gorohovski, A., Mukherjee, S. & Frenkel-Morgenstern, M. Non-optimal codon usage preferences of coronaviruses determine their promiscuity for infecting multiple hosts. The FEBS journal 288, 5201–5223, doi:10.1111/febs.15835 (2021).

33 Butt, A. M., Nasrullah, I., Qamar, R. & Tong, Y. Evolution of codon usage in Zika virus genomes is host and vector specific. Emerging microbes & infections 5, e107, doi:10.1038/emi.2016.106 (2016).

34 Wu, X. et al. Optimization and deoptimization of codons in SARS-CoV-2 and the implications for vaccine development. bioRxiv, doi:10.1101/2022.09.03.506470 (2022).

35 Goodman, L. B. et al. A point mutation in a herpesvirus polymerase determines neuropathogenicity. Plos Pathog 3, 1583–1592, doi:10.1371/journal.ppat.0030160 (2007).

36 Duraffour, S. et al. Mutations Conferring Resistance to Viral DNA Polymerase Inhibitors in Camelpox Virus Give Different Drug-Susceptibility Profiles in Vaccinia Virus. J Virol 86, 7310–7325, doi:10.1128/Jvi.00355-12 (2012).

37 Tang, X. et al. Evolutionary analysis and lineage designation of SARS-CoV-2 genomes. Science Bulletin 66, 2297–2311, doi:10.1016/j.scib.2021.02.012 (2021).

38 Tang, X. et al. On the origin and continuing evolution of SARS-CoV-2. National Science Review 7, 1012–1023, doi:10.1093/nsr/nwaa036 (2020).

39 Qian, Z., Li, P., Tang, X. & Lu, J. Evolutionary dynamics of the severe acute respiratory syndrome coronavirus 2 genomes. Medical Review, 3–22, doi:10.1515/mr-2021-0035 (2022).

40 Yan, X., Hoek, T. A., Vale, R. D. & Tanenbaum, M. E. Dynamics of Translation of Single mRNA Molecules In Vivo. Cell 165, 976–989, doi:10.1016/j.cell.2016.04.034 (2016).

41 Presnyak, V. et al. Codon optimality is a major determinant of mRNA stability. Cell 160, 1111–1124, doi:10.1016/j.cell.2015.02.029 (2015).

42 Buhr, F. et al. Synonymous Codons Direct Cotranslational Folding toward Different Protein Conformations. Mol Cell 61, 341–351, doi:10.1016/j.molcel.2016.01.008 (2016).

43 Yu, C. H. et al. Codon Usage Influences the Local Rate of Translation Elongation to Regulate Co-translational Protein Folding. Mol Cell 59, 744–754, doi:10.1016/j.molcel.2015.07.018 (2015).

44 Chen, F. et al. Dissimilation of synonymous codon usage bias in virus-host coevolution due to translational selection. Nat Ecol Evol 4, 589–600, doi:10.1038/s41559-020-1124-7 (2020).

45 Poland, G. A., Kennedy, R. B. & Tosh, P. K. Prevention of monkeypox with vaccines: a rapid review. The Lancet Infectious Diseases 22, e349–e358, doi:10.1016/S1473-3099(22)00574-6 (2022).

46 Aksamentov, I., Roemer, C., Hodcroft, E. & Neher, R. Nextclade: clade assignment, mutation calling and quality control for viral genomes. Journal of Open Source Software 6, 3773, doi:10.21105/joss.03773 (2021).

47 Cingolani, P. et al. A program for annotating and predicting the effects of single nucleotide polymorphisms, SnpEff: SNPs in the genome of Drosophila melanogaster strain w1118; iso-2; iso-3. Fly (Austin) 6, 80–92, doi:10.4161/fly.19695 (2012).

48 Yang, Z. PAML 4: phylogenetic analysis by maximum likelihood. Mol Biol Evol 24, 1586–1591, doi:10.1093/molbev/msm088 (2007).

